# Pattern reinstatement and attentional control: overlapping processes during episodic long-term memory retrieval

**DOI:** 10.1101/2022.02.21.481285

**Authors:** Melinda Sabo, Daniel Schneider

**Author notes:** Address of correspondence: Melinda Sabo, Leibniz Research Centre for Working Environment and Human Factors Ardeystraße 67 44139 Dortmund Germany.

## Abstract

Episodic long-term memory (eLTM) retrieval involves the reinstatement of neural patterns from the encoding phase. However, recent evidence suggests that comparable effects on cortical activity patterns can also be linked to attentional control processes on the level of memory representations. The current investigation assesses these two processes independently based on alpha-beta-band activity in the electroencephalogram (EEG). During encoding, subjects were presented with an object on a certain position on the screen and had to imagine it on a new position. In each trial, either the task-irrelevant presentation position or the task-relevant imagination position was lateralized. In the retrieval phase, subjects first made an old/new judgement based on centrally presented objects and then reported the imagination position. Pattern reinstatement should be reflected in similar lateralized alpha-beta activity during encoding and retrieval. Conversely, the influence of attentional control processes during retrieval would be associated with the suppression of alpha-beta power contralateral to the to-be-reported imagination position and with the increase of activity contralateral to the irrelevant presentation position. Our results support this latter pattern. This shows that an experimental differentiation between selective attention and pattern reinstatement processes is necessary when studying the neural basis of eLTM retrieval on cortical level.

## Introduction

The uniqueness of episodic memory consists in its capacity to allow individuals to consciously re-experience past events (i.e., to do mental time travel) by retrieving details of a particular episode^1^. One central mechanism determining episodic memory retrieval is ecphory. The term was defined by Tulving as a synergistic process through which information stored in the form of memory traces interacts with the retrieval cue to facilitate the recollective experience^2^. Thus, in Tulving’s view, the engram of an event alone is not sufficient unless it is accompanied by a compatible retrieval cue^2^, which prompts a rapid, unconscious reactivation of sensory memory traces^3, 4^. Moreover, the retrieval cue seems to not only trigger an early information reactivation, but also the replay of neural patterns from the encoding phase.

Since Tulving’s phenomenological description of ecphory, several studies aimed to explore this phenomenon at a neurophysiological level. The results of these studies suggest that episodic memory retrieval entails a very early “replay” or reinstatement of cortical patterns present during the original event^5, 6^, which can happen as early as 100-300 ms after cue presentation^7^. This finding proves to be robust across studies using different brain imaging techniques (e.g., functional magnetic resonance imaging; fMRI^8,9,10^, EEG^7^, intracranial EEG^11^, magnetoencephalogram^12^) and different types of to-be-remembered stimuli (e.g., words and imagined objects^13^, verbal recall of short movies^5^). Moreover, disruption of this early cortical reinstatement process is suggested to have detrimental effects on episodic memory performance, thus showing that early replay is functionally relevant for episodic memory retrieval^4^.

Yet, a recent study conducted by Sutterer and colleagues^14^ questions the idea that cortical activity underlying pattern reinstatement reflects a single process. In their paper, the authors suggest that alpha oscillations (8-12 Hz) code for spatial locations associated with different objects in the study phase and these patterns re-emerge in the retrieval phase.

Nonetheless, they further argue that this activity might as well reflect sustained attention to spatial locations linked to each object. Similarly, Waldhauser and colleagues^4^ reveal that lateralized alpha-beta oscillations track spatial positions in an analogous way during encoding and later retrieval. The authors interpret their results as evidence for pattern reinstatement. However, one cannot exclude the possibility that the underlying alpha activity reflects an overlap between pattern reinstatement and further processes, such as attentional control mechanisms.

A considerable amount of research demonstrates the involvement of alpha oscillations in tracking the focus of attention, both at a perceptual level^15–17^ and within working memory representations^18–22^. The latter finding is especially evident in paradigms, in which the relevance of different items held in working memory changed throughout the task, requiring subjects to switch the focus of attention from one item representation to another. This process of prioritization can, for example, be tracked by lateralized alpha oscillations, showing a contralateral decrease with respect to currently relevant item representations and contralateral increase corresponding to irrelevant ones^23, 24^. Consequently, lateralized alpha activity is not only suggested to play an important role in deploying attention to relevant memory representations, but also in inhibiting irrelevant ones^25–28^. The idea of attentional mechanisms playing an active role in eLTM retrieval is further supported by studies aiming to understand the involvement of the posterior parietal cortex (PPC) in episodic memory retrieval. For instance, it is suggested that activity observed during episodic memory retrieval in the two sub-regions of the PPC, the medial intraparietal sulcus (IPS) and superior parietal lobule, reflects deployment of top-down attention^29^. Similarly, a recent review highlights the importance of the lateral IPS and postcentral gyrus in the manipulation and transformation of information according to the current task goals, a processes argued to occur rather late^30^ and highly likely as the result of attentional mechanisms^29^. These conclusions are also in line with the predictions of the Attention to Memory (AtoM) model^31^, which emphasizes the role of bottom-up and top-down attention in episodic memory retrieval^31^. Accordingly, bottom-up attention captures goal relevant information once this is retrieved by the medial temporal lobe (MTL), while top-down attention is important for maintaining the retrieval goal and for guiding the search for relevant information. The second mechanism becomes especially decisive in cases of poor memories, requiring more effortful memory searches^30, 32^.

Taken together, a considerable amount of research confirms the necessity of pattern reinstatement for episodic memory retrieval^7–12^. However, there is also evidence that attention facilitates the retrieval of information from episodic memory^29–31^. As such, it seems plausible that the cortical activity thought to be a marker of pattern reinstatement reflects some further operations, potentially overlapping during retrieval. In this context, it becomes highly unclear whether task-specific modulations of cortical activity previously reported during eLTM retrieval (e.g., alpha oscillations^4, 14^) are a neural correlate of pattern reinstatement or also of attentional control processes during eLTM retrieval. We therefore designed an EEG experiment to answer this question and to shed light on the exact role of changes of cortical activity (here measured by topographic changes of alpha and beta oscillations) during eLTM retrieval.

The experiment consisted of three phases: an encoding phase, a distractor phase (involving counting backwards in steps of three), and finally a retrieval phase. In the encoding phase, subjects were presented with an object, which could appear on four positions: left, right, top or bottom. Afterwards, the object disappeared from the screen and participants’ task was to imagine it on a new position as indicated by a cue presented prior to the object (showing +90° or -90°). Finally, the retrieval phase started with the presentation of old or new objects appearing in the center of the screen. Participants were first required to make an old/new judgement, followed by a display indicating to report the position where the previously seen object had been imagined. The experiment was designed in a way that each trial associated two positions to the objects: a presentation position and an imagination position. The position, where the object had been presented did not contain specific task-relevant information and was thus not required for task completion. The imagination position was task-relevant, and it had to be reported during the retrieval phase. Moreover, the imagination position was always located 90° clockwise or counterclockwise compared to the original presentation position, leading to the association of a lateral (left vs. right) and a central position (top vs. bottom) to each object (figure 1).

**Figure 1.**
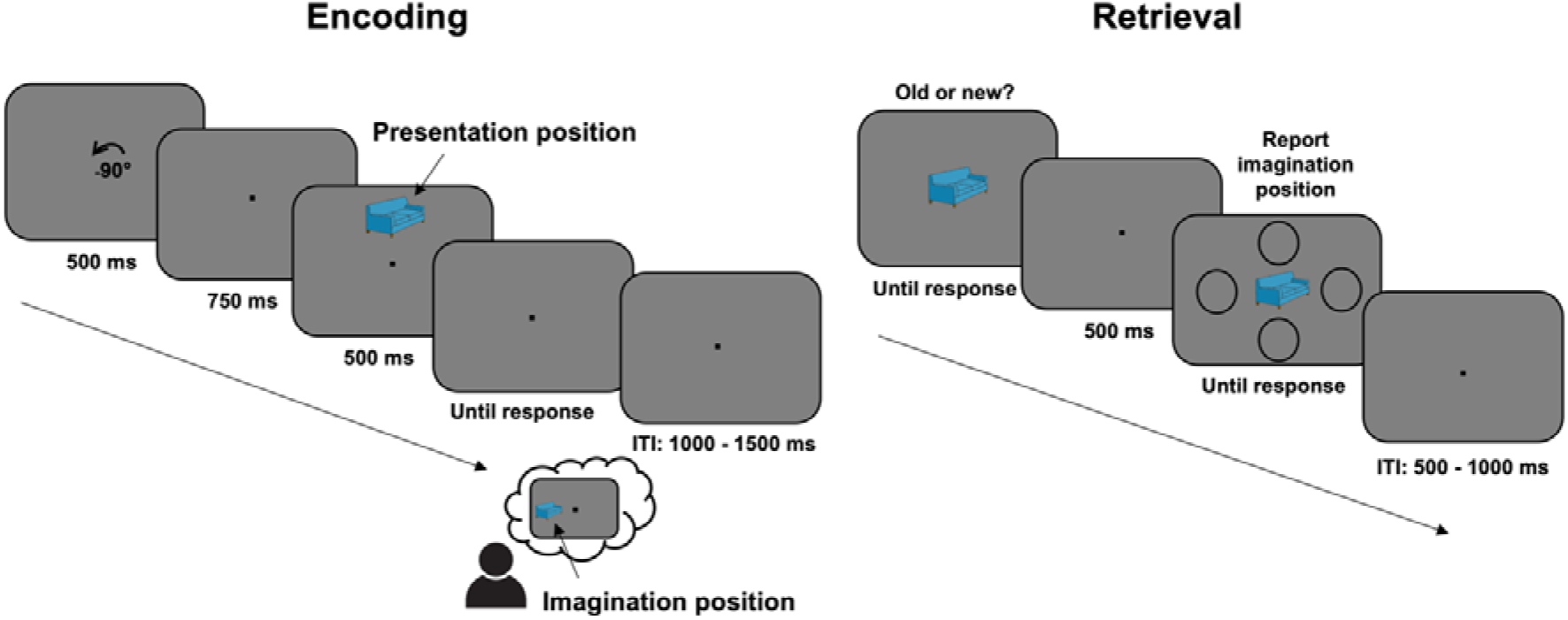
Schematic illustration of the experimental procedure. In the encoding phase, subjects were required to learn associations between everyday objects and imagination positions (top, bottom, left or right). Each object was presented three times during the encoding phase and was supposed to be associated with the same imagination position. In the retrieval phase, subjects completed first an old/new recognition task, in which objects from the previous phase were randomly interleaved with a set of 120 previously unseen objects. If participants reported being familiar with the respective item, they were asked to report the associated imagination position. At the end of each trial, an inter-trial interval (ITI) randomly varying between 1,000-1,500 ms (encoding phase) or 500-1,000 ms (retrieval phase) was introduced.

The experiment was designed in a way that for each object the lateral position was either task-relevant (when being the imagination position) or task-irrelevant (when being the presentation position), which enabled us to independently measure the lateralization of alpha-beta power (8-20 Hz) in each experimental condition. If the modulation of alpha-beta oscillations as a function of the location associated with the individual object does indeed reflect spatial attentional processes, we expect to find a contralateral alpha-beta decrease (relative to the ipsilateral hemisphere), linked to the selective retrieval of the imagination position^23–25, 27^. Importantly, in line with prior research on the attentional selection of working memory content, this effect should be accompanied by a contralateral alpha-beta power increase related to the inhibition of the irrelevant presentation position^25–28^. Moreover, we assume that these effects are already evident during the retrieval of episodic information for the old vs. new decision (i.e., prior to the display requiring the imagination position report) (see figure 2). However, if the modulation of alpha-beta power as a function of the location associated with the retrieved information rather reflects reinstated oscillatory patterns from the encoding phase^4^, we should be able to observe a stronger alpha-beta decrease contralateral (as compared to ipsilateral) to both the presentation and the imagination position (as both effects are expected to be present during information encoding) (see figure 2). Although subjects are not required to explicitly memorize the presentation position, this information is processed together with the imagination position during encoding and potentially retrieved later. Overall, the current study is suitable to provide experimental information on the extent to which oscillatory patterns measured by EEG/MEG are appropriate for representing ecphoric processes during eLTM retrieval.

**Figure 2.**
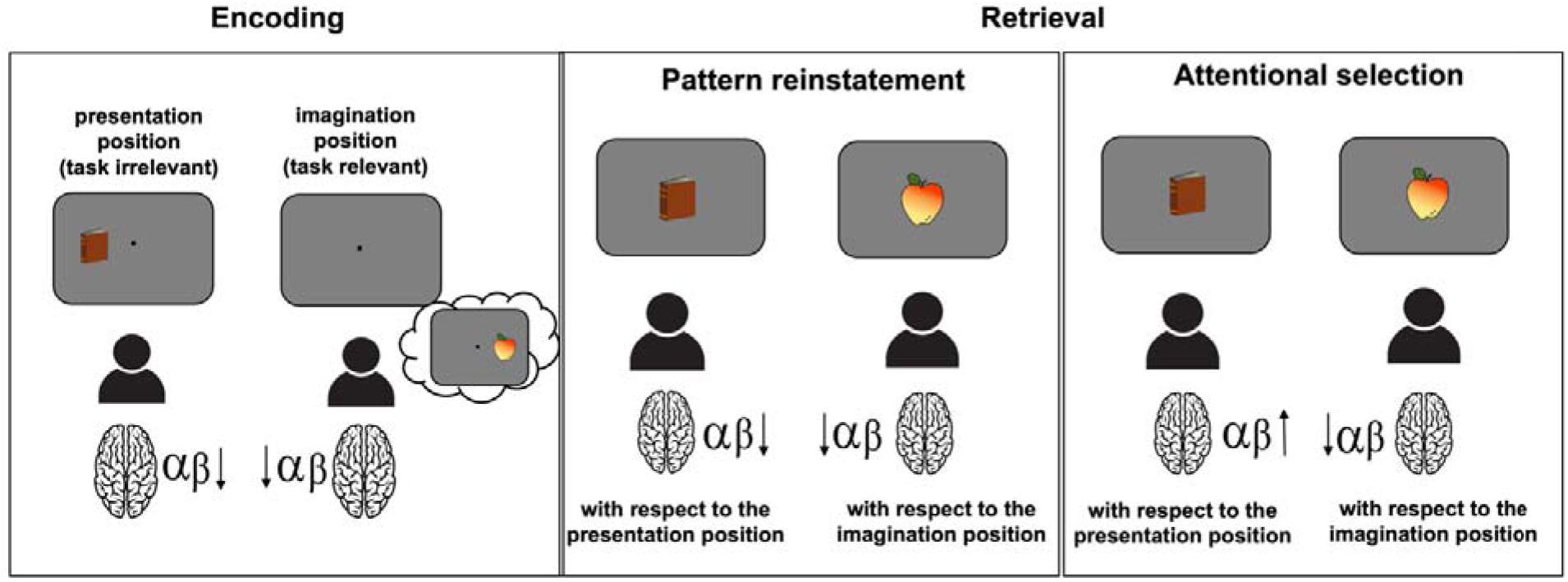
Schematic illustration of the hypotheses. Based on previous findings^17, 18^, we expect to find a contralateral alpha-beta decrease with respect to both the presentation position, as well as to the imagination position in the encoding phase. As for the retrieval phase, two alternative hypotheses were formulated. If lateralized alpha-beta-band activity reflects pattern reinstatement, a contralateral alpha-beta decrease is expected both with respect to the presentation and to the imagination position. However, if alpha-beta oscillations reflect attentional selection mechanisms, a contralateral alpha-beta increase with respect to the task-irrelevant information and a decrease with respect to the task-relevant information is expected.

## Results

### Behavioral Data

Figure 3a depicts the accuracy distribution (in percent) for both the old/new recognition task (*M* = 91.06, *SD* = 7.12), as well as for the imagination position recall (*M* = 81.13, *SD* = 15.18). In both cases, subjects had an above chance performance: *t*(41) = 37.34, *p* < .001, 95% CI [88.84, 93.28] (old/new recognition task), *t*(41) = 23.96, *p* < .001, 95% CI [76.40, 85.86] (position recall task). Figure 3b shows the reaction time distribution for the two tasks. Moreover, we tested whether any of the incorrect positions was more frequently reported, by comparing the number of reports of each incorrect position. The repeated-measure ANOVA revealed a main effect of irrelevant position. Since sphericity was violated (χ^2^(41) = 10.71, *p* = .004, ε = 0.80), Greenhouse-Geisser corrected results are reported: *F*(1, 41) = 10.33, *p* < .001, η_p_ ^2^ = 0.20. The subsequent post-hoc results revealed that participants reported the presentation position (*M* = 7.71, *SD* = 6.14) more frequently, as compared to the other two incorrect positions (*M_position1_* = 4.90, *SD_position1_* = 4.28; *M_position2_* = 5.00, *SD_position2_* = 4.74): *t*(41) = 3.81, *p_adj_* = .001, *d_av_* = 0.53, 95% CI [1.32, 4.29] (compared to incorrect position1) and *t*(41) = 3.31, *p_adj_* = .002, *d_av_* = 0.49, 95% CI [1.06, 4.36] (compared to incorrect position2). There was no difference between the frequency of reports for the remaining two incorrect positions: *t*(41) = -0.18, *p_adj_* = .85, *d_av_* = 0.02, 95% CI [-1.13, 0.94] (figure 3c).

**Figure 3.**
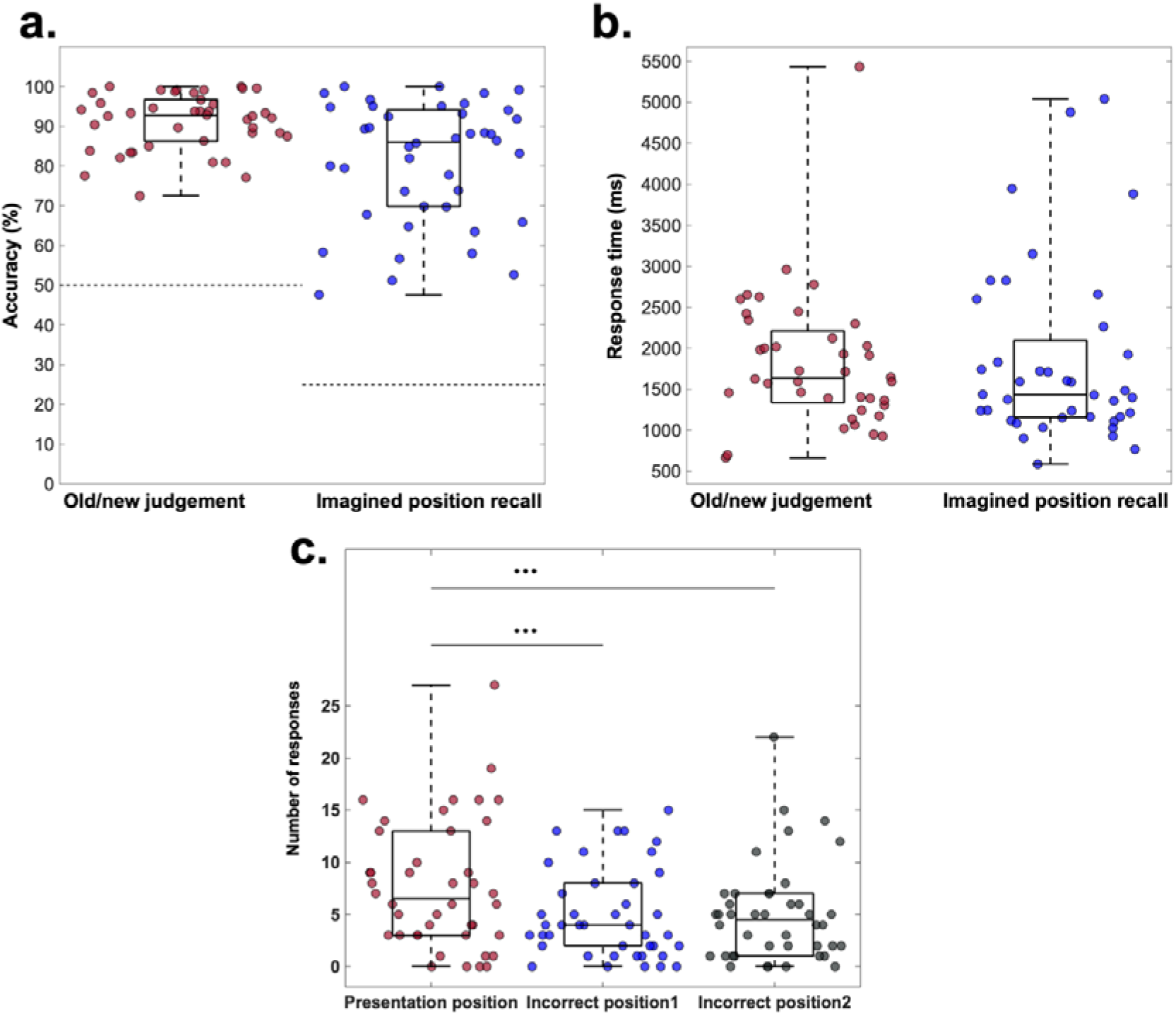
Behavioral performance. a. Boxplot illustrating the distribution of average accuracy (%) for the old/new recognition and imagination position recall task. All participants show above-chance performance (chance level - recognition task: 50%; recall task: 25%). **b.** Boxplot showing the distribution of average reaction times (ms) for the two tasks: *M_recognition_* = 1538.80 ms, *SD_recognition_* = 102.70 ms; *M_recall_* = 1,364.10 ms, *SD_recall_* = 199.25 ms. **c.** Boxplot illustrating the distribution of incorrect responses. The Y-axis denotes the frequency of reporting the respective incorrect position, where the maximum values is 30.

### Lateralized alpha-beta power oscillations

Figure 4a depicts the results of the cluster-based permutation analysis (for details see Methods), in which contralateral and ipsilateral alpha-beta (8-20 Hz) power was compared within each condition (i.e., relative to the presentation- and to the imagination position) in the encoding phase. Results indicated a significant cluster between 101-817 ms after stimulus onset with respect to the presentation position, while no significant cluster was found in the comparison relative to the imagination position (see in addition figure 5a and 6a for the topographical and time-frequency representation of the encoding phase data).

**Figure 4.**
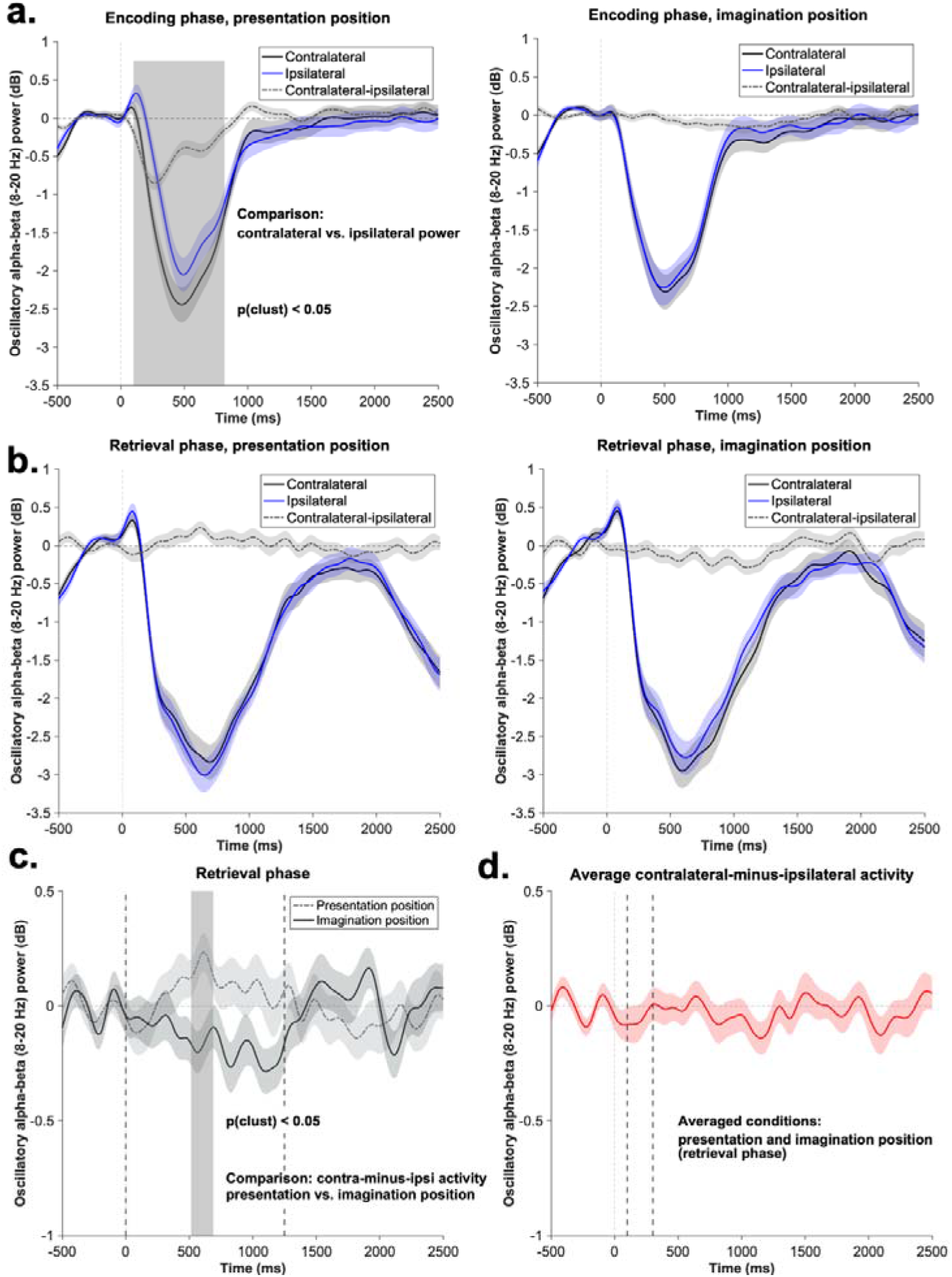
Lateralized posterior alpha-beta analysis. Data were obtained from a posterior electrode cluster: TP7/8, P5/6, P7/8, PO7/8. The shaded area represents the standard error of the mean activity. **a.** Time course of the contralateral and ipsilateral alpha-beta power (8-20 Hz) during the encoding phase, with respect to presentation position (left) and imagination postion (right). The gray area marks the significant cluster (*p* < .05) obtained as a results of the cluster-based permutation procedure used to contrast the contralateral an ipsilateral activity (time window: 0-2,582 ms). **b.** Time course of the contralateral and ipsilateral alpha-beta power (8-20 Hz) during retrieval phase, with respect to the presentation position (left) and the imagination postion (right). The contralateral and ipsilateral activity was contrasted using a cluster-based permutation approach (time-window: 0-2,582 ms). **c.** Comparison of the contralateral-minus-ipsilateral alpha-beta power between the two conditions of the retrieval phase (i.e., relative to the presentation and imagination positions). The gray area marks the significant cluster (*p* < .05) obtained as a results of the cluster-based permutation procedure. The procedure was conducted on the 0-1,250 ms time window, as indicated by the gray dotted lines. **d.** Time course of the contralateral-minus-ipsilateral power averaged for the two conditions of the retrieval phase.

**Figure 5.**
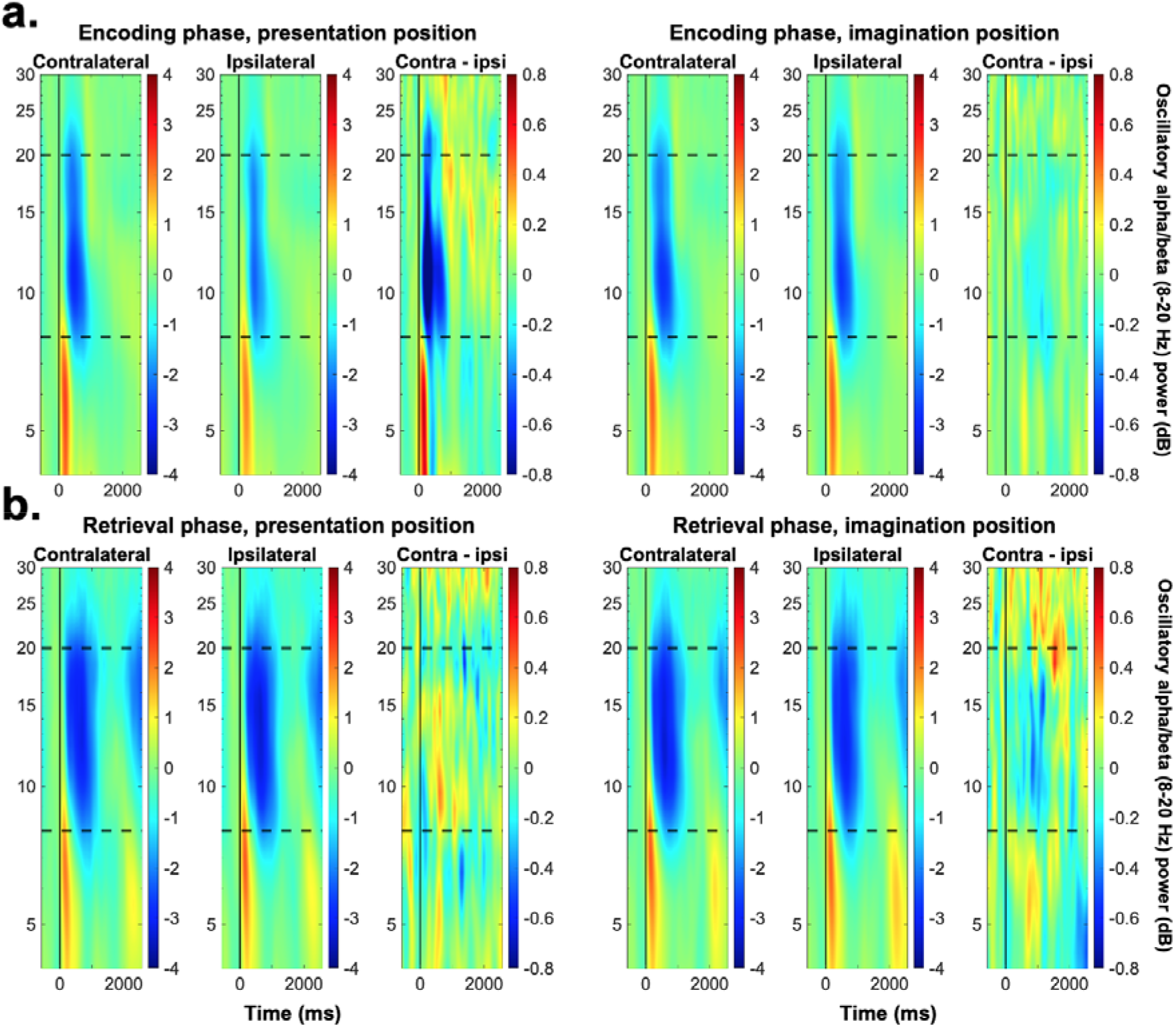
Time-frequency representation of the contralateral, ipsilateral, and contralateral-minus-ipsilateral data. The activity-of-intereset was obtained from a posterior electrode cluster: TP7/8, P5/6, P7/8, PO7/8. Dotted lines indicate the frequencies of interest (8-20 Hz).

**Figure 6.**
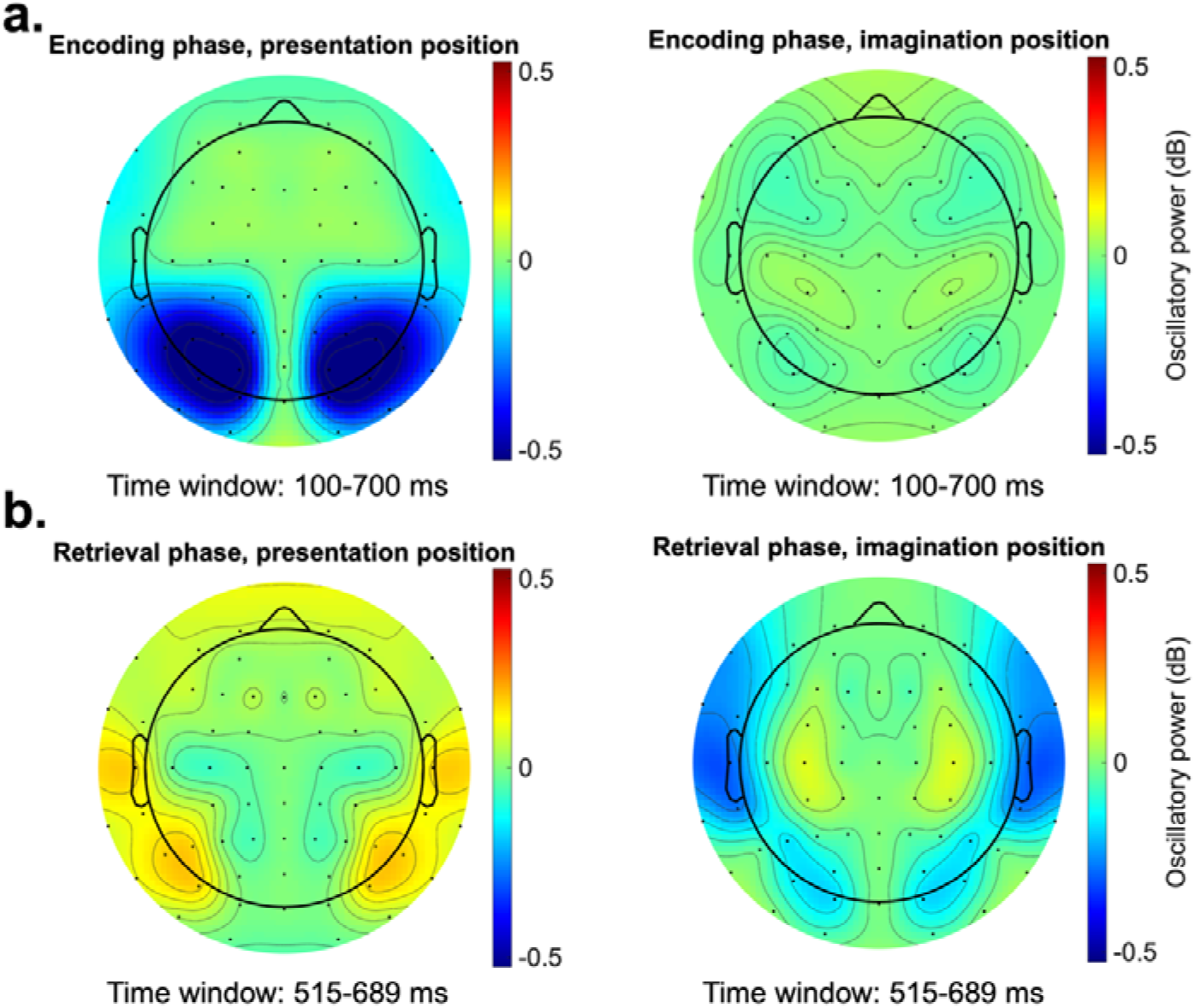
Topographical distribution of the contralateral-minus-ipsilateral alpha activity. The activity-of- intereset was obtained from a posterior electrode cluster: TP7/8, P5/6, P7/8, PO7/8.

For the retrieval phase analysis, two competing hypotheses were tested (see figure 2). As shown in figure 4b, no significant clusters were found when contrasting the ipsilateral and contralateral activity either relative to the presentation- or to the imagination position. In order to bring further evidence in support for these results, a cluster-based permutation procedure was applied to test whether the contralateral-minus-ipsilateral activity averaged for the two conditions in the retrieval phase significantly differs from zero. This analysis yielded no significant clusters (figure 4d). The subsequent one-sample t-test (see Methods) further confirms these results. As such, we found that the activity averaged over the 100-300 ms time window (*M* = -0.05, *SD* = 0.43) did not differ from zero: *t*(41) = -0.82, *p* = .41, 95% CI [-0.19, 0.07]. Overall, this suggests that no evidence was found in support for the pattern reinstatement account.

For the second analysis (testing the attentional selection account), we contrasted the contralateral-minus-ipsilateral activity between the two conditions (i.e., the condition, in which the activity was relative to the imagination position and the one relative to the presentation position). The results of the cluster-based permutation analysis revealed a significant cluster between 515-689 ms (figure 4c). Importantly, in the subsequent post-hoc analyses (see Methods), data from these significant time windows were averaged and tested against zero in one-tailed one-sample t-tests for each condition (see Methods for further details). Results showed that the contralateral-minus-ipsilateral activity significantly differed from zero in both conditions (presentation condition: *M* = 0.18, *SD* = 0.45; imagination condition: *M* = -0.15, *SD* = 0.53). First, we observed an alpha-beta decrease contralateral to the imagination position, suggesting that subjects selectively focused attention on the task relevant position during retrieval, *t*(41) = -1.91, *p_adj_* = .03, 95% CI [-Inf -0.01]. Second, an alpha-beta increase was found contralateral to the presentation position, indicating that subjects inhibited the task irrelevant position, *t*(41) = 2.61, *p_adj_* = .01, 95% CI [0.06, Inf].

These results thus indicate that lateralized alpha-beta oscillations present during eLTM retrieval reflect attentional control processes (see in addition figure 5a and 6a for the topographical and time-frequency representation of the retrieval phase data).

### Brain-behavior correlation

In order to assess whether the frequency of reporting the presentation position is linked to the oscillatory brain patterns, we correlated the presentation position ratio with the contralateral-minus-ipsilateral activity measured with respect to the presentation position (figure 7a, for more details see Methods). Since this analysis was exploratory, correlations were calculated for the entire time window and across all frequencies ranging from 4 to 30 Hz. In order to account for multiple comparisons, a cluster-based permutation analysis was conducted for the time window of interest (0-1250 ms, as indicated in the Methods section). This analysis revealed a significant cluster in the lower alpha (∼8-10 Hz) frequency range between 0-600 ms after stimulus onset, depicting a negative correlation between the presentation position ratio and the amplitude of the contralateral-minus-ipsilateral difference with respect to the presentation position (figure 7b). That is, the higher the contralateral vs. ipsilateral increase in alpha power relative to the presentation position, the lower the frequency of reporting the presentation position (compared to the total amount of errors made). This result could potentially suggest that subjects, who systematically reported the presentation position, had difficulties inhibiting this spatial information.

**Figure 7.**
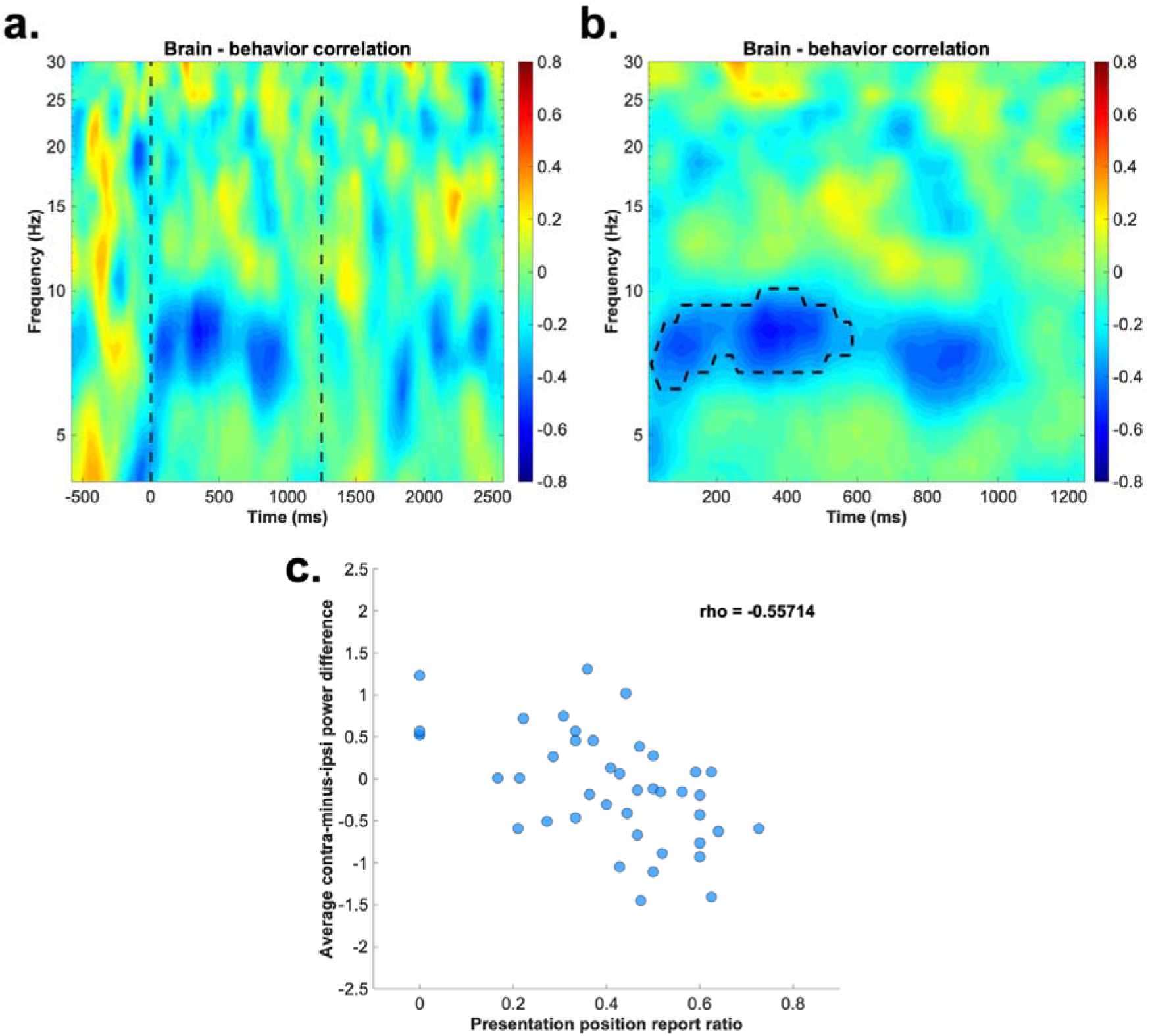
Results of brain-behavior correlations. a. Spearman’s correlation coefficient depicting the correlation between the contralateral-minus-ipsilateral activity with respect to the presentation position and the presentation position ratio across the whole epoch and all frequencies. The dotted line indicates the time window used for the cluster-based permutation analysis (0-1,250 ms). **b.** Results of the cluster-based permutation procedure. Dotted line indicates the significant cluster (*p* < .05). **c.** Scatterplot of the negative correlation between the two parameters of interest: presentation position ratio and the contralateral-minus-ipsilateral activity averaged over the significant cluster (shown in panel b). Please note that this figure is shown only for illustrative purposes.

## Discussion

The aim of the current study was to investigate whether similarities in cortical activity during the encoding and retrieval of long-term episodic memories qualify as a clear marker of pattern reinstatement during eLTM retrieval. We addressed this question by measuring topographic changes of the oscillatory power in the alpha and beta frequency range during the encoding and retrieval of objects and associated spatial positions. As such, we designed an eLTM experiment, in which each object was associated with a task-relevant imagination position and a task-irrelevant presentation position, one of which was lateralized in each trial. Based on previous evidence, we formulated two alternative hypotheses. Accordingly, the pattern reinstatement account predicts that activity patterns during encoding and retrieval should be similar. More specifically, if lateralized alpha-beta oscillations represent a marker of pattern reinstatement, we expected to observe a stronger alpha-beta decrease contralateral (compared to ipsilateral) to both the presentation and imagination position, as both positions had to be processed during the encoding phase. Conversely, if alpha-beta-band activity during eLTM retrieval tracks attentional control mechanisms, a stronger alpha-beta power decrease contralateral (as compared to ipsilateral) to the task-relevant imagination position and a reversed effect with respect to the presentation position (i.e., a contralaterally increased alpha-beta power) were expected during retrieval.

At a behavioral level, results indicated that subjects reported the presentation position more systematically in comparison with the other incorrect positions. This clearly supports the idea that the presentation position was encoded together with the object, despite being task-irrelevant for the position recall. This is not surprising, since a considerable amount of literature argues about the special role of spatial location in organizing information in memory^33–35^. In line with this, prior research suggested that even spatial information associated with only passively viewed stimuli is encoded into eLTM and can be retrieved in a later recall task^4^. The phenomenon we observed at the behavioral level is analogous to that of committing swap errors in working memory paradigms (i.e., erroneous report of task-irrelevant working memory content)^36^. It has been suggested that these swap errors occur with higher frequency as the memory load increases^37, 38^ and are possibly due to the confusion of the target feature with highly similar non-target information^39^. In the context of the current experiment, it seems possible that the remaining residual information of the task-irrelevant presentation position interfered with the target report, thus leading to a more systematic bias towards the presentation position, especially in cases of poor memories.

The lateralized alpha-beta analysis conducted for the encoding phase revealed a strong contralateral decrease with respect to the presentation position (relative to the ipsilateral oscillatory power), suggesting that subjects moved the focus of attention to the object’s position^15–17^. This was expected, since the visual objects were presented unilaterally (i.e., they drew attention in a bottom-up way^4^). A slight alpha-beta power asymmetry was also found with respect to the imagination position but did not reach significance in the current sample. Nevertheless, saccades were observed during the imagination task (see supplementary material), indicating a directional biasing of gaze towards the imagination position. Although subjects were specifically instructed to fixate on the centrally presented point during the imagination task, their gaze involuntarily moved towards the location, where the object had to be imagined. This phenomenon is consistent with findings indicating that saccades and microsaccades (i.e., small saccades happening during fixation) occur during attentional orienting to perceptual stimuli^40^, as well as during attentional selection of relevant memory representations^41, 42^. As such, based on this evidence, it seems plausible that the saccades we observed during the imagination task reflected a process of attentional re-orienting to the imagination position.

Lateralized alpha-beta activity was also observed during the retrieval phase. The contralateral-minus-ipsilateral power differed between the presentation and imagination conditions, showing a strong contralateral-minus-ipsilateral suppression relative to the task-relevant imagination position and a reversed effect relative to the task-irrelevant presentation position. This suggests that the two lateralized effects are linked to separate mechanisms, both facilitating eLTM retrieval. According to prior research on the attentional selection of working memory contents^23, 25, 27^, this pattern of results can be considered as evidence for attentional re-focusing on the task relevant imagination position (reflected by the contralateral alpha-beta power decrease) and the inhibition of the task-irrelevant presentation position (reflected by the contralateral alpha-beta power increase). As indicated by the behavioral results, it seems plausible that the presentation position was also encoded together with the imagination position into eLTM, despite being task irrelevant. As such, the centrally presented objects in the retrieval phase (acting as a retrieval cue) triggered the reactivation of the associated imagination position, which was accompanied in certain trials by an automatic reactivation of the presentation position, a process presumably interfering with the retrieval of the task-relevant spatial information. In this context, it seems plausible that the interference was reduced by attentional control processes, required for inhibiting irrelevant information and for selecting the relevant spatial position for the later report. The correlational analysis offered further evidence in favor of this explanation. Results suggested that the lower the amplitude of the contralateral alpha-beta increase with respect to the presentation position, the greater the probability of erroneously reporting it. In agreement with our interpretation, this would suggest that subjects, who had difficulties inhibiting the irrelevant information, were more likely to be biased towards reporting the presentation position.

The pattern of results obtained in the current study corroborate the findings provided by experiments investigating attentional control on the level of working memory.

Accordingly, it was argued that attentional control is involved in monitoring the attentional switches between memory representations, a process which is tracked by lateralized alpha activity. More specifically, task-relevant items entering the focus of attention were shown to elicit a stronger contralateral alpha suppression, compared to the ipsilateral hemisphere, while the currently irrelevant items are related to a stronger contralateral alpha increase^25, 26, 43^.

Considering the similarity of these findings with the current result patterns, it seems plausible that upon object presentation, subjects retrieved both spatial associations (i.e., presentation and imagination positions) from long-term memory. Subsequently, in support for the position report, this information was brought back into working memory, where the task-relevant position was selected, while the task-irrelevant one was inhibited. In this context, it seems likely that the attentional selection mechanisms acting at the level of working memory facilitated the process of long-term memory retrieval.

This interpretation is also in line with an influential model of memory (i.e., Embedded process model of working memory), which states that successful information retrieval from eLTM is realized by passing the respective information through the focus of attention in working memory^44^. In accordance with this theory, Fukuda and colleagues^45^ provided experimental evidence suggesting that information retrieved from long-term memory is brought back into working memory, where it is temporarily stored. As such, the authors discuss about the existence of two different inter-connected memory states relevant for the retrieval process: an active state of information representation in working memory and a passive state of information storage in long-term memory. Importantly, a recent study conducted by Vo and colleagues^46^ further supports the idea of working memory and long-term memory retrieval being connected. In their study, authors demonstrated that activity in the occipital (V1-V4 areas) and parietal cortex (V3AB, IPS0-2 retinotopic regions) during working memory maintenance and long-term memory retrieval is highly similar^46^. Moreover, this conclusion is fully in line with the results of an earlier study, in which a classifier trained on the cortical activity recorded during a long-term memory task performed above chance when tested on the delay period of a working memory task^47^. Authors interpreted their results as evidence for the long-term memory system supporting information maintenance in working memory.

The present results have important implications for the study of eLTM retrieval. First, we provide experimental evidence supporting the active role of selective attention during eLTM retrieval, a process, which can be easily confounded with pattern reinstatement especially when assessing neural correlates of retrieval based on cortical activity. Therefore, we question whether brain modulations recorded on the scalp surface by EEG/MEG techniques constitute a suitable method for the investigation of pattern reinstatement.

Moreover, previous evidence shows that hippocampal pattern completion and MTL activity mediate the process of pattern reinstatement^11, 48^. As such, it seems plausible that the hippocampus and some parts of the MTL are necessary for cortical reinstatement. In line with these findings and the results of the current study, we argue that future studies aiming to investigate the phenomenon of pattern reinstatement during eLTM should carefully consider two important aspects: (i) the choice of the brain imaging technique; and (ii) the constructions of experiments, which allow for an adequate control of overlapping processes, such as selective attention.

## Conclusions

To summarize, the current study aimed to shed light on the functional role of analogous activity patterns during eLTM encoding and retrieval. Namely, we addressed the question of whether these modulations reflect pattern reinstatement or attentional control mechanisms. Our results supported the latter account, showing that subjects selected the task relevant information, while they inhibited the task irrelevant presentation position. These results raise an important issue for the study of eLTM retrieval: there is a need to differentiate and control for the effects of attentional selection processes, especially when investigating the neural correlates of pattern reinstatement.

## Methods

### Participants

Originally, data from 46 participants was collected. Four participants had to be excluded from the final analysis: three of them were not able to complete the task using the given visual strategy and in the case of the fourth one, technical issues with the EEG system were encountered. The remaining 42 participants (22 female, 20 male) had an age between 18 and 30 years (*M* = 24.1 years, *SD* = 2.83), were right-handed, had normal or corrected-to-normal vision, and did not suffer from any neurological or psychiatric disorders. For their participation, subjects received a compensation either as a payment of 10 €/hour, or study participation credits (required for the completion of a bachelor’s degree in psychology). Prior to participation, written consent was obtained from all subjects. The study was in accordance with the Declaration of Helsinki and had been approved by the ethics committee of the Leibniz Research Centre for Working Environment and Human Factors (Dortmund, Germany).

### Procedure

Upon arrival, participants were presented with general information about the study, followed by the completion of the German version of the following questionnaires: a demographic survey and the Edinburgh Handedness Inventory^49^. As a next step, the EEG cap was set and subjects were led into the sound-attenuated, electrically shielded, and dimly lit EEG laboratory.

The experiment was performed on a 22-inch CRT monitor (100 Hz; 1024 x 768 pixels), with a viewing distance of approximately 145 cm. The task was realized in Lazarus IDE (Free Pascal) and presented using ViSaGe MKII Stimulus Generator (Cambridge Research Systems, Rochester, UK). The eLTM task included a short training and three phases: encoding, distractor, and retrieval phase. Once the task was finalized, subjects completed a follow-up questionnaire, in which they were asked about the strategy they used and difficulties they encountered while completing the experiment. The whole session lasted between 3 and 3.5 hours.

### Stimulus material

A total number of 240 everyday objects were used for the eLTM task^50^. Since the original object pool consisted of 260 images, a prior image selection process was conducted. First, images depicting objects pointing towards certain directions (e.g., a gun, a pointing finger) were excluded (seven objects in total). Secondly, we conducted a survey, in which an independent sample of subjects rated the remaining 253 objects based on a set of criteria (see details in the next section). Based on these results, 13 more objects were removed from the final pool of images.

### Image rating survey

The image rating survey contained three statements, each reflecting the agreement of subjects with respect to: (i) proper luminance, contrast and vividness of the object; (ii) ease of recognizability; (iii) complexity. Subjects’ task was to rate all objects based on these three statements. Answers were given on a 5-point Likert scale (for statements one and two: 1 = “Total disagreement”, 5 = “Total agreement”; for statement three: 1 = “Simple”, 5 = “Complex”). The aim of the image rating survey was to identify the objects, which stood out in terms of sensory quality (reflected by luminance, contrast, and vividness) and those, which were easily recognizable. In addition, we aimed to obtain measures of complexity, which were later used to counterbalance the stimuli in the eLTM task.

A total number of 30 participants completed the questionnaire (18 females, 10 males, and 2 unspecified gender). Their age ranged between 18 and 35 years (*M* = 27 years, *SD* = 4.38). The survey was realized in PsychoPy3^51^ and uploaded online to Pavlovia (https://pavlovia.org/). The questionnaire was completed by volunteers, who did not receive compensation for their participation.

In order to identify the 13 to-be-removed objects, a score reflecting the mean object quality (statement one) and recognizability (statement two) was calculated (*M* = 4.57, *SD* = 0.25). The objects obtaining the 13 lowest scores (smaller than 4.07) were excluded from the stimulus pool. Furthermore, for categorizing objects based on complexity, we conducted a median split analysis on the remaining 240 images, resulting in a pool of 120 simple and 120 complex objects.

### Episodic long-term memory task

The experimental task started with a training, which contained 27 encoding trials and 19 retrieval trials and aimed to familiarize subjects with the experimental procedure. Thus, data acquired during these practice trials was excluded from the final analysis. Once the training was complete, subjects performed the encoding phase. Each encoding trial started with a cue, presented on a gray background (RGB: 128, 128, 128) depicting either +90° (i.e., the imagination position is situated clockwise relative to the presentation position) or -90° (i.e., the imagination position is situated counterclockwise relative to the presentation position). Following a 750 ms inter-stimulus interval (ISI), objects (size: 4.3° x 3°) were presented on one of the four positions on the screen (top, bottom, left, right). The distance between the object and the central fixation point was 3° visual angle. After 500 ms, the object disappeared and the subject’s task was to imagine the object on a new position, as indicated by the cue shown at the beginning of the trial. For instance, if the cue showed +90° and the object was presented at the top position, participants had to imagine the object on the right position. Both during object presentation and the imagination task, participants were instructed to fixate on the centrally presented black dot (size: 0.2° x 0.2°). The imagination task was self-paced. After each trial subjects had to confirm through a mouse click that they finished imagining the object on the corresponding position on the screen. The inter-trial interval (ITI) varied randomly between 1000 and 1500 ms (figure 1).

The encoding phase consisted of 360 trials, organized in six blocks (60 trials/block). Between blocks, self-paced breaks were introduced. Each participant learnt 120 object- imagination position associations. In order to facilitate the learning process, each object occurred in three trials, each time on the same position and having the same imagination position associated with. However, trials showing the same objects could not be in the same block. The object-imagination position associations were randomized across subjects, while object complexity and format (landscape vs. portrait) were counterbalanced across the imagination condition.

The encoding phase was followed by a distractor phase, which required participants to count backward starting from 500, in steps of three. The distractor phase had a duration of three minutes and was preceded and followed by three-minute breaks. Finally, in the retrieval phase, subjects were shown a centrally presented object (0.2° x 0.2°) and their task was to decide whether they had already seen the respective item (old/new judgement). Half of the items presented in this phase were also shown in the encoding phase, whereas the rest of them were unknown to participants. Responses were given with the left and right button of the computer mouse each button being associated with a response type (old or new). The mapping between the button and the response was counterbalanced across subjects. In trials, in which subjects indicated familiarity with the object, the associated imagination position had to be further reported by clicking on a circle (diameter: 2°) associated with the remembered position. The distance between the central object and each of the four circles was 3° visual angle. Trials in the retrieval phase were separated by an ITI of 500-1000 ms. The retrieval phase consisted of 240 trials in total (120 old and 120 new items), organized in 4 blocks which were separated by self-paced breaks (figure 1).

### Data Analysis

All analyses were conducted in MATLAB® (R2019b).

### Behavioral analysis

As a first step in the behavioral analysis, we tested whether participants’ performance (as reflected by the average accuracy) in the old/new recognition and the imagination position recall task were above chance. In both cases, we conducted a separate one-sample t-test, in which we contrasted the average accuracy to 50% (old/new judgement) and 25% (imagination position recall). Furthermore, in order to investigate whether subjects reported more frequently any of the incorrect positions, the number of reports corresponding to each incorrect position was compared using a one-way analysis of variance (ANOVA). Only trials, in which subjects provided both a familiarity judgement (i.e., reported whether the object was old or new) and a valid imagination position report (i.e., clicked within the circle representing the imagination position) were included in the behavioral analysis. Failure of clicking within the indicated circle led to trials with missing responses, which were excluded from further analysis.

### EEG recording and preprocessing

EEG data were recorded using a 64 Ag/AgCl passive electrode system (Easycap GmbH, Herrsching, Germany). The electrodes were distributed according to the extended 10/20 System^52^ and the signal was amplified by a NeuroOne Tesla AC-amplifier (Bittium Biosignals Ltd, Kuopio, Finland). Data were sampled at a rate of 1000 Hz and a 250 Hz low-pass filter was applied during recording. AFz served as ground electrode, FCz as reference electrode. Throughout the recording session, impedances were kept below 20 kΩ . For the EEG data analysis, we used the EEGLAB toolbox^53^ (v. 14.1.2) implemented in MATLAB® (R2019b). Before preprocessing, each subject’s dataset was separated into an encoding and a retrieval dataset, so subsequent analyses were done separately on the encoding and retrieval data. Moreover, in order to speed up the analysis of the retrieval dataset, we only included trials showing objects present during the encoding phase (i.e., old objects).

As a first preprocessing step, data was 0.5 Hz high-pass (filter order: 6600, transition band-width: 0.5 Hz, cutoff frequency at -6dB: 0.25 Hz) and 30 Hz low-pass filtered (filter order: 440, transition band-width: 7.5 Hz, cutoff frequency at -6dB: 33.75 Hz) using a Hamming windowed sinc FIR filter (i.e., pop_eegfiltnew from the EEGLAB toolbox). Next, using the automated channel rejection procedure implemented in the toolbox (i.e., pop_rejchan), artifact channels were rejected (range: 0 – 6, Encoding dataset: *M* = 2.33 channels, Retrieval dataset: *M* = 2.04 channels), with the use of the following parameters: kurtosis of each channel as a rejection measure, outlier selection based on 15 standard deviations. Next, the data were re-referenced to the average activity of the remaining electrodes.

In order to prepare the data for the independent component analysis (ICA), the data were further downsampled to 200 Hz, 1 Hz low-pass filtered (filter order: 660, transition band-width: 1 Hz, cutoff frequency at -6dB: 0.50 Hz) using a Hamming windowed sinc FIR filter and epoched. For both the encoding and the retrieval dataset, the chosen time window was -1000-3000 ms, time-locked to the object’s onset. Afterwards, the data were baselined to the 200 ms time window prior to stimulus onset and an automated trial rejection procedure implemented in EEGLAB (i.e., pop_autorej; threshold: 500 μV, maximum % of rejected trials: 5%) was used to reject trials containing extreme fluctuations (encoding phase: *M* = 50.04 trials, 15.35%; retrieval phase: *M* = 8.16 trials, 10.85%).

Following, ICA was conducted on the rank reduced data (remaining number of channels minus one). Dimensionality reduction was achieved using the principal component analysis implemented in EEGLAB. Next, the obtained IC weights were transferred to the bandpass-filtered, average re-referenced data. In order to identify independent components (IC) reflecting blinks, horizontal - and vertical eye movements and general discontinuities, we used the ADJUST plugin^54^ (version 1.1.1). In addition, in order to identify the ICs containing high residual variance, we estimated a single-equivalent current dipole model, using the DIPFIT plug-in. Thus, further components exceeding 40% residual variance regarding their dipole solution were excluded. Overall, 15.95 IC components were marked for exclusion in the encoding (range: 6-35) and 15.42 in the retrieval dataset (range: 3-25). Once data was epoched and baseline-corrected (same parameters as above), the marked IC components were removed. Afterwards, trials left with extreme fluctuations were excluded from the analysis using the automated approach mentioned above (threshold: 1000 μV, maximum % of rejected trials: 5%): on average 74.40 trials (20.50%) were excluded from the encoding dataset and 12.90 trials (14.26%) from the retrieval dataset. Finally, channels excluded at the beginning of the analysis were interpolated using a spherical spline interpolation technique.

### Time-frequency decomposition

Time-frequency decomposition was done by convolving each trial of the EEG data with 3-cycle complex Morlet wavelets. A complex Morlet wavelet is defined as a complex sine wave that is tapered by a Gaussian. The obtained epochs contained 200 time points in the time window -582-2582 ms relative to the object onset (for both leaning and retrieval phase) and the frequencies ranged between 4 and 30 Hz, increasing in logarithmic steps of 26. The number of cycles, which defines the width of the tapering Gaussian, increased linearly as a function of frequency by a factor of 0.5. As such, the number of cycles for the lowest frequency (4 Hz) was three, whereas 11.25 cycles were used for the highest frequency (30 Hz). A time window between -500-0 ms was chosen as a spectral baseline. The time-frequency decomposition was conducted through the STUDY functions of EEGLAB (i.e., std_precomp). For this analysis, all encoding phase trials remaining after the trial rejection procedure (conducted during preprocessing) were included in the STUDY. In case of the retrieval phase data, we additionally excluded the trials, in which subjects failed to correctly identify the old objects. Moreover, trials in which subjects’ reaction time in the recognition task was greater than 3 s were also excluded.

### Alpha-beta power lateralization

In order to answer our main research question, lateralized modulations of posterior alpha-beta (8-20 Hz) power were assessed^4^. As a first step in our analysis, we focused on the activity patterns of the encoding phase. This activity was obtained by averaging the power values across the frequency band of interest (8-20 Hz) and across a lateral posterior electrode cluster, consisting of PO7/8, P7/8, P5/6, TP7/8. The electrode selection procedure was based on previous findings on attentional selection within working memory^20, 27, 55–57^. The mean contralateral and ipsilateral power was separately calculated with respect to the presentation and imagination position. To assess the difference between the contralateral and ipsilateral activity within each condition (i.e., relative to the presentation position and to the imagination position), we applied a cluster-based permutation approach for each condition separately. The procedure was conducted on the time window 0-2582 ms. First, the power value of the contralateral and ipsilateral activity was compared at each of the 163 time points using paired-sample t-tests. This resulted in a vector of 163 *p*-values corresponding to each comparison of the original data. Based on this vector, clusters containing more than one significant *p*-value (i.e., lower than 0.05) were obtained. In the following step, we generated a distribution of maximum cluster sizes. This was achieved by randomly re-shuffling the condition labels (i.e., contralateral and ipsilateral) 1000 times and in each iteration paired-sample t-tests were used to compare the ipsilateral and contralateral activity at all time points. Based on the vector of 163 p-values, the cluster containing the largest number of significant *p*-values (i.e., smaller than 0.05) was subtracted and saved for each iteration, thus resulting in a distribution of maximum cluster sizes. Finally, we determined the 95^th^ percentile of the cluster size distribution and clusters bigger than this cutoff value were considered significant.

In order to address the two competing hypotheses regarding the alpha-beta oscillatory patterns during the retrieval phase (see figure 2), two separate analyses were conducted. First, the pattern reinstatement account predicts a contralateral alpha-beta decrease with respect to both the presentation and imagination positions (see figure 2 middle). To test this assumption, we calculated the mean contralateral and ipsilateral activity relative to the initial presentation and imagination position. Similar to the encoding phase analysis, data were obtained from the lateral posterior electrode cluster (PO7/8, P7/8, P5/6, TP7/8), by averaging the activity across the 8-20 Hz frequency band. Following, we calculated the contralateral-minus-ipsilateral activity and averaged it across the two relevant conditions (i.e., with respect to the presentation and imagination position). Subsequently, the average power values were contrasted to zero using the same cluster-based permutation procedure described above (time window: 0-2582 ms). Then, using a one-sample t-test, we tested whether the same activity averaged over the time window 100-300 ms significantly differs from zero. The choice of the time window was motivated by previous EEG/MEG studies, which showed that cortical patterns from the encoding phase are reinstated during memory retrieval ∼100-300 ms after object presentation^4, 7, 58^. Furthermore, using the same cluster-based permutation approach described above, we contrasted the contralateral and ipsilateral activity (time window: 0-2582 ms) within each condition (i.e., relative to the presentation and to the imagination position).

Second, we tested the attentional selection account, according to which we expected: (i) a stronger contralateral (as compared to ipsilateral) alpha-beta power decrease relative to the imagination position; and (ii) a stronger contralateral (as compared to ipsilateral) increase relative to the presentation position (see figure 2 right). To determine a time window during which these lateralized effects could be tested, the contralateral-minus-ipsilateral difference waves obtained with respect to the imagination and presentation position were contrasted using the same cluster-based permutation approach (see above). This procedure was restricted to the time window from 0 to 1250 ms (after object presentation), during which effects of selective attention usually appear in the EEG correlates^59, 60^. In case of a significant effect, we conducted *t*-tests against zero based on the activity averaged over the significant cluster, with the aim of testing whether a contralateral suppression (imagination position) or a contralateral increase in alpha power (presentation position) were evident.

### Brain-behavior correlations

Beyond the lateralized alpha-beta analyses, a further exploratory correlational analysis was conducted in order to investigate whether a systematic report of the presentation position (in case of an erroneous position report) is related to the modulations of lateralized alpha-beta oscillations. As a behavioral parameter, we calculated a so-called “presentation position ratio”, obtained by dividing the number of presentation position reports with the total number of incorrect responses. Regarding the EEG parameter, we considered the contralateral-minus-ipsilateral power values relative to the presentation position in the retrieval phase. Since no previous study investigated this effect during eLTM retrieval, we did not have a hypothesis about which time windows or frequencies to use in this analysis. Consequently, we calculated the Spearman correlations between the power value at each time (-582-2582 ms) x frequency (4-30 Hz) combination and the behavioral parameter. In order to adjust for multiple comparisons, a cluster-based permutation procedure (see above) was applied on all time-frequency data points in the time-window from 0-1250 ms (see time window selection procedure above).

### Inferential statistics and effect sizes

All statistical tests and measures of effect size were obtained in MATLAB® (R2019b). For the one-way ANOVA comparing the frequency of report for each incorrect position, Mauchly’s test for sphericity was applied. Because the sphericity assumption was violated, the Greenhouse-Geisser correction was introduced (denoted with ε). As a measure of effect size, we calculated partial eta squared (η_p_ ^2^). Regarding the applied t-tests, two-tailed *p*-values were calculated, if not otherwise specified. As a measure of effect size, we followed the recommendations of Lakens^61^ and computed Cohen’s *d_av_*. In the case of post-hoc tests requiring a correction for multiple comparison, false discovery rate (FDR) corrected p-values were provided (denoted by *p_adj_)* ^62^.

## Acknowledgements

The authors would like to thank Tobias Blanke for programming the experiment and Pia Deltenre, Barbara Foschi, and all associated student assistants for their help in the data collection.

## Author contribution

M.S. was responsible for data collection. M.S. and D.S. designed the experiment, conducted the analyses and wrote the manuscript. D.S. acquired the necessary funding.

## Data availability statement

The data and all the scripts used for the reported analyses are publicly available on Open Science Framework (OSF) accessing the following link: https://osf.io/ngdzm/?view_only=018451896c84482e8df189cee11646a8.

## Supplementary material

**Figure 1.**
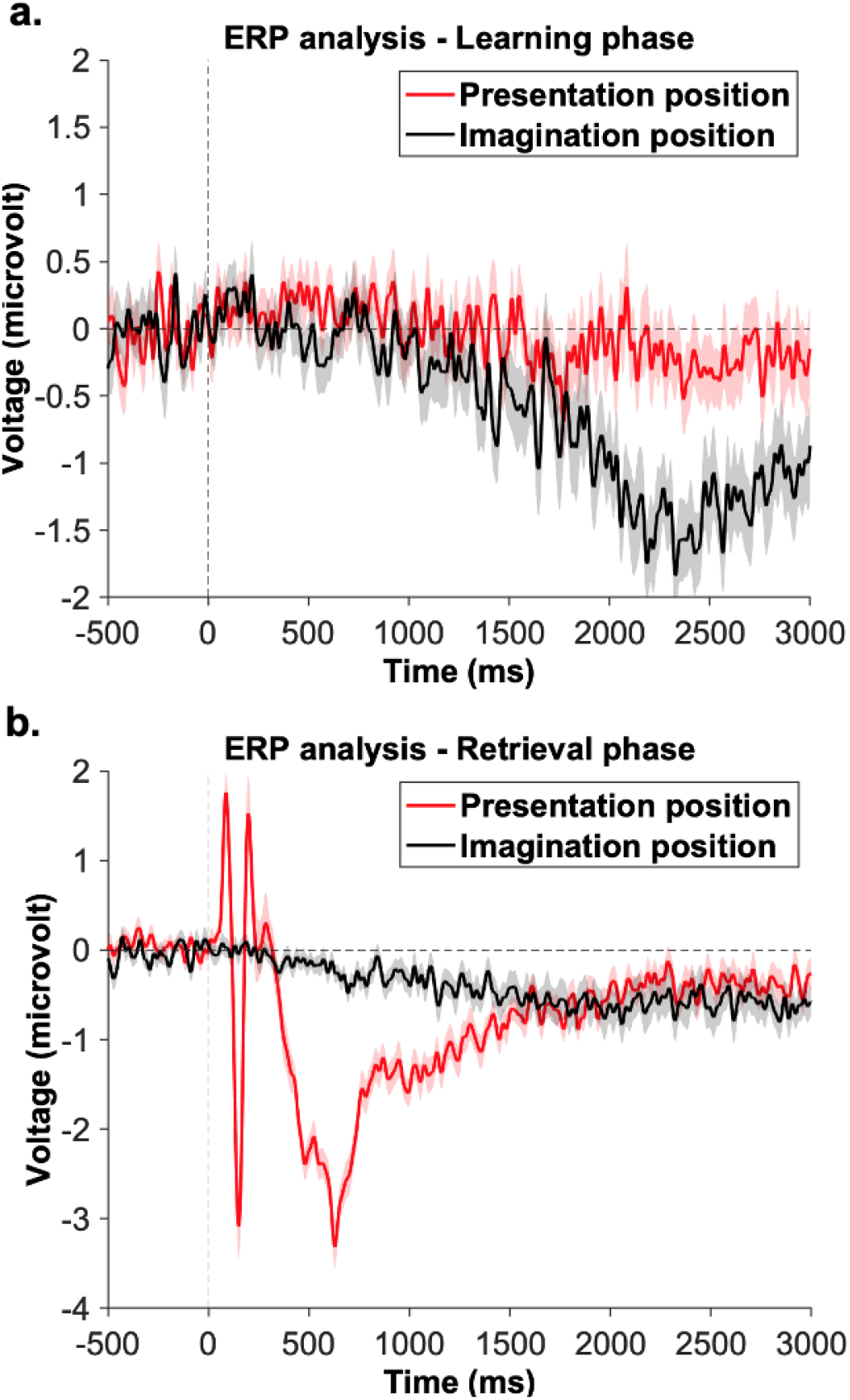
Time course of the event-related potentials. The figure depicts the contralateral-minus-ipsilateral activity relative to the presentation and imagination positions. Similar to the lateralized alpha-beta analyses, data were obtained by averaging the contralateral and ipsilateral activity from the lateralized posterior electrode cluster (TP7/8, PO7/8, P5/6, P7/8). The data used for the current analysis includes the independent components containing lateral eye movement, which were not excluded in the artifact removal procedure (via ADJUST during preprocessing). The shaded area represents the standard error of the mean activity. **a.** Comparison of contralateral-minus-ipsilateral ERPs relative to presentation and the imagination position of the learning phase. Clear saccadic activity can be observed with respect to the imagination position. **b.** Comparison of contralateral-minus-ipsilateral ERPs relative to presentation and the imagination position of the retrieval phase. This shows that saccadic activity does not overlap with the time window of the observed alpha-beta effect.

